# High coverage genome sequencing and identification of genomic variants in Bengal tiger (*Panthera tigris tigris*)

**DOI:** 10.1101/306399

**Authors:** P Anuradha Reddy, Harish Kothandaraman, P V Parvati Sai Arun, Anil Challagandla, Dushyant Singh Baghel

**Affiliations:** CSIR-Centre for Cellular and Molecular Biology, Uppal Road, Hyderabad, India-500007; Nucleome Informatics Private Limited, Genome Block, Miyapur, Hyderabad, India-500049

## Abstract

Bengal tiger (*Panthera tigris tigris*), one of six extant tiger subspecies, occurs solely in the Indian subcontinent. Although endangered and threatened by various extinction risks, this is the most populous tiger subspecies with the highest genetic diversity and strongest chance of survival in the wild. Availability of high quality genomic information on this animal will help us understand its ability to adapt to different habitats and environmental changes, in addition to comparative studies with other subspecies. Here we report high coverage sequencing of the Bengal tiger genome and its mapping to the Amur tiger genome in order to discover single nucleotide to large structural variants. A total of 345 Gb, roughly equivalent to 144X coverage of the genome, was generated from 1,149,381,669 raw read pairs. Further, 990,060,729 clean read pairs, again equivalent to 115X coverage, were retained from the raw read data and considered for comparative analysis with the Amur tiger genome. This alignment showed that 97.35% of the bases mapped at 5X depth, 97.26% at 10X and 90.44% at 50X depth. We identified a total of 3,601,882 single nucleotide variants, 948 structural variants, 56,649 copy number variants and 1,760,347 simple sequence repeats. We report the first high coverage genome sequence of Bengal tiger with an overview of its genomic variants when compared to the Amur tiger genome. Of the several variants identified, we further have to assess and validate variants potentially associated with the ability of the animal to adapt to environmental changes, disease susceptibility and other important biological phenomena.

## Introduction

Over the last few decades, application of genetics to conservation research and management has greatly contributed to our understanding of the dynamics of endangered populations, with precise empirical data on genetic diversity, population structure, gene flow and effective population size, amongst various demographic parameters [1, 2]. Exponential advances in next-generation sequencing technologies (NGS) and reduced sequencing costs have further enhanced our ability to cost-effectively sequence complete genomes of several species. However this deluge of genomic data, although cost-effective, only marginally improves the accuracy of analysis and our perception of various population demographic parameters described above. On the other hand, as discussed by McMahon et al. [2], large scale genomic data can help us identify regions in the genome important for evolutionary processes such as speciation, adaptation to local conditions thereby providing clues about the species’ ability to survive in a changing world [3–7]. As we sequence high coverage genomes of individuals, we will be able to discover single nucleotide variations to large structural variants which are associated with local adaptations or disease susceptibility.

In this paper we report high coverage (144X raw data and 115X clean data) sequencing of the Bengal tiger (*Panthera tigris tigris*) genome and its mapping to the Amur tiger (*Panthera tigris altaica*) genome [8]. Interestingly these two extant tiger subspecies occur in extremely diverse habitats and environmental conditions. Amur tiger occurs exclusively in sub-temperate, snow covered habitats. Bengal tiger, on the other hand, occupies diverse tropical habitats varying from Himalayan foothills and Gangetic floodplains, mangrove forests of Sunderbans, Central India plateau, and Western Ghats. Although India today has the largest number of tigers, approximately 2200 [9], these are spread out across several small tenuously connected populations, each with questionable long-term survivability in isolation. Worryingly, the increasing genetic drift caused by isolation of meta-populations may actually interfere with genetic signatures of local adaptations thereby challenging the species’ viability.

For a very long time it was believed that single nucleotide variants (SNVs) contribute to a majority of the individual’s genomic variations [10, 11]. Now it is recognised, albeit poorly understood, that much larger changes in the genome like large structural variants (SVs) and copy number variants (CNVs) also contribute significantly to diseases [12–16], phenotypic variations [17], and immunity [18, 19]. Discovery and research on structural variants are largely in humans [20], and to some extent in domestic and semi-wild animals [21–25]. To the best of our knowledge, this is the first report on the discovery of copy number variants and large structural variants in the genome of a wild, endangered species. This comprehensive data on simple sequence repeats (SSRs), SNVs, SVs and CNVs discovered through comparative analysis of a high coverage Bengal tiger genome sequence with that of the Amur tiger will initiate our understanding on genomic changes and the species’ ability to adapt to discreet habitats.

## Data Description

### Sample collection

We isolated DNA from post-mortemly collected kidney tissue of an approximately nine month old male tiger cub. The cub, named Dhanush (ID No. M01157), was rescued from Metiguppe Forest Division, Karnataka on 15^th^ January 2015 and housed at Sri Chamarajendra Zoological Gardens, Mysuru till its death possibly due to pneumonia on 2^nd^ February 2015 (Letter from Assistant Director, SCZG, Mysuru, dated 03/02/2015). DNA was isolated with DNeasy Blood and Tissue kit (Qiagen, Germany) according to manufacturer’s instructions. Approximately 5μg of purified DNA was used for library preparation.

### Library Preparation and Sequencing

We prepared multiple paired-end libraries (insert sizes of 250 bp, 350 bp and 550 bp) and mate-pair libraries (insert sizes of 5 kb and 10 kb) with Truseq Nano DNA HT sample preparation kit FC-121-4003 as per the manufacturer’s protocols. We purified the generated PCR products on AMPure XP system and selected required sizes on Agilent 2100 Bioanalyzer. We ran the final selected libraries of each insert size in multiple lanes on Illumina HiSeq 2500 with 2×150 chemistry till optimal coverage for all analyses was obtained. A total of 345 Gb was generated from 1,149,381,669 raw read pairs from all paired-end and mate-pair libraries (Supplementary Information Table S1). This is roughly equivalent to 144X coverage of the genome based on an estimated genome size of 2.4 Gb [8].

### Data Cleaning and Quality Control

We cleaned the library data to remove adaptors, trim low quality bases and also to remove ‘N’ with Trimmomatic v0.36 [26]. In case of mate-pair libraries, the generated data was also checked for presence and removal of Cre-LoxP adaptors. After cleaning the data and quality control assessment we had an average of 86.3% clean data from all libraries. A total of 94.35% reads cleared Q20 while 86.67% reads had Q30 quality (Table 1, Supplementary Information Table S2). A total of 990,060,729 clean read pairs were retained from the entire raw read data and considered for further analysis (Supplementary Information Table S1). This again is roughly equivalent to 115X coverage of the genome. We further checked this clean data for contamination by homology searches of 10,000 reads from each lane, from different insert size libraries, against the NCBI Nt database with BlastN [27]. BlastN searches resulted in homology hits centred around *Felis catus* (domestic cat), *Panthera tigris* (tiger), *Neofelis nebulosa* (clouded leopard).

**Table 1:**
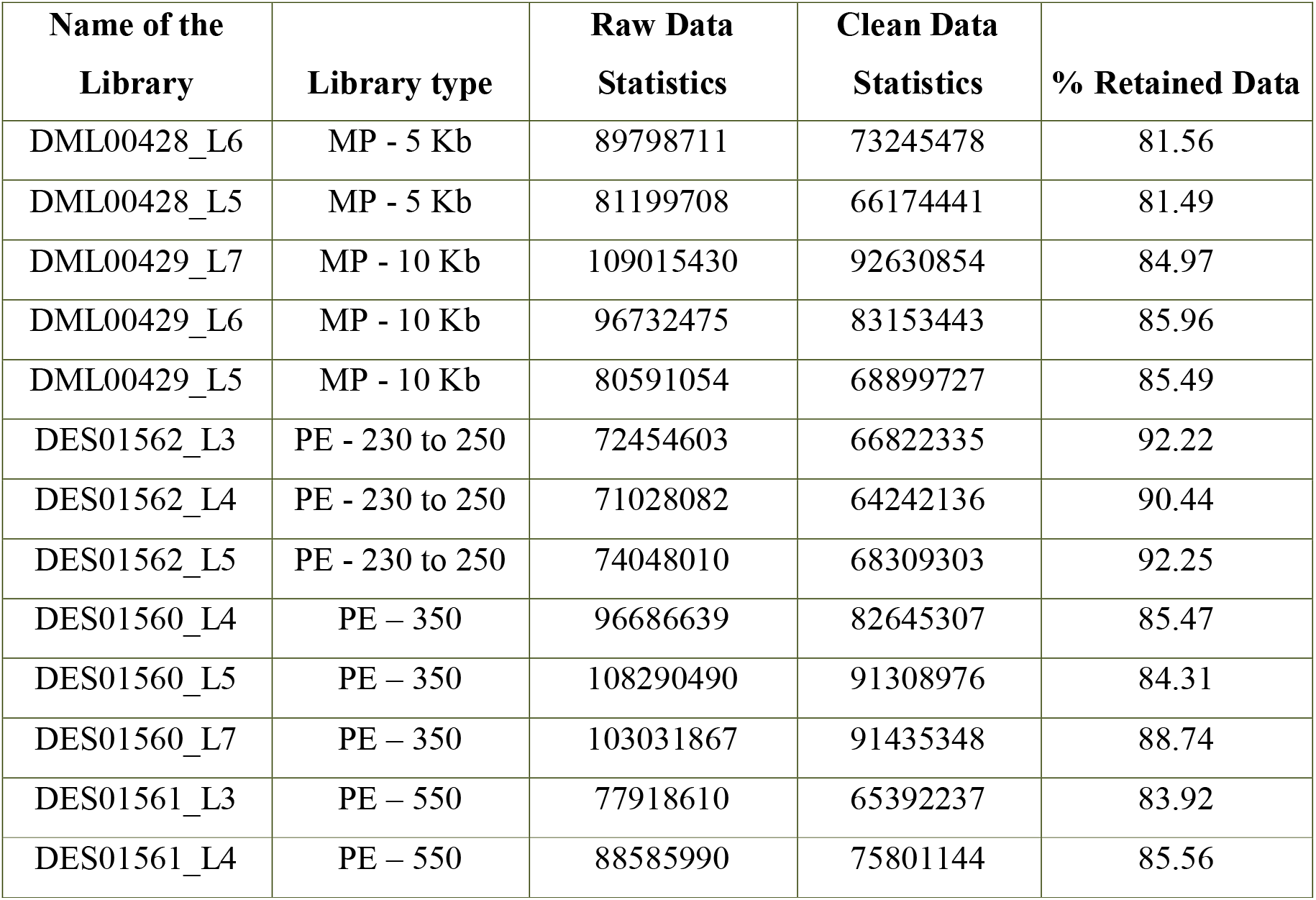
Summary statistics of multiple mate pair (MP) and pair-ended (PE) library sequencing data. Percentage of retained data was calculated as: % Retained Data = (Clean data statistics of each library /Raw data statistics of each library) *100

### Mapping to the Amur Tiger Genome

We used the Amur tiger genome [8] as a reference for this study. We mapped our clean reads to the Amur tiger genome with MEM module of BWA (Burrows-Wheeler Aligner) with default parameters [28]. This alignment showed that 97.35% of the bases mapped at 5X depth, 97.26% at 10X and 90.44% at 50X depth (based on de-duplication statistics) (Supplementary Information Table S1). Picard Tools from Broad Institute were used to add read groups, and to sort and mark bam files duplicates generated from the alignment of sequences to the reference genome (https://broadinstitute.github.io/picard/). Clean reads of multiple inserts were combined using Samtools [29]. We used HaplotypeCaller module of Genome analysis toolkit (Broad institute) to identify variants in the merged bam files (https://software.broadinstitute.org/gatk/documentation/tooldocs/3.80/org_broadinstitute_gat_k_tools_walkers_haplotypecaller_HaplotypeCaller.php). Insertions and deletions >60 bp were treated as structural variants (SVs) and were identified with Breakdancer-max [30]. We also used Lumpy SV with stringent filters to identify SVs (https://github.com/arq5x/lumpy-sv). Single nucleotide variants (SNVs) identified by Gatk were hard-filtered with Bcftools (http://www.htslib.org/doc/bcftools-1.0.html) based on MQ60, DP >10 and Quality >90. The identified SNVs were annotated with snpEff tool (http://snpeff.sourceforge.net/). To identify copy-number variants (CNVs), we used CNVnator (Abyzov) (https://github.com/abyzovlab/CNVnator) at a bin size of 200. In order to identify simple sequence repeats (SSRs), the predicted SNVs and InDels were plugged back into the reference genome to get a consensus sequence of the Bengal tiger genome. This consensus was then used to identify SSRs using MISA (http://pgrc.ipk-gatersleben.de/misa/). Breakdancer generated a total of 27,758 SVs with a minimum of three reads to support each variant, while Lumpy SV generated 948 SVs with a minimum of 100 reads to support each variant. These SVs were further classified into intra-chromosomal translocations, inversions, deletions and insertions (Table 2).

**Table 2:** Structural variants identified in the Bengal tiger genome and their counts. BND - chromosomal translocations; INV - inversion; DEL - deletion; DUP - duplication.

| Types of Structural variants | Counts |
| --- | --- |
| BND | 730 |
| INV | 0 |
| DEL | 191 |
| DUP | 27 |
| **Total** | **948** |

Similarly, we identified 3,601,882 SNVs in the Bengal tiger genome when compared to Amur tiger genome (Table 3).

**Table 3:** Statistics describing the SNVs identified in the Bengal tiger genome. SNPs - Single nucleotide polymorphisms; INS - Insertions; DEL - Deletions.

| Types of Single nucleotide variants | Counts |
| --- | --- |
| SNPs | 3007160 |
| INS | 298801 |
| DEL | 295921 |
| **Total** | **3601882** |

While analysing copy number variants with CNVnator, we identified 56,649 CNVs such as deletions or duplications, based on the insert-size metrics at various loci (Table 4).

**Table 4:** Copy number variants identified in the Bengal tiger genome, their types and their respective counts.

| Types of Copy number variants | Counts |
| --- | --- |
| Deletion | 44910 |
| Duplication | 11739 |
| **Total** | **56649** |

Using MISA (http://pgrc.ipk-gatersleben.de/misa/), we identified a total of 1,760,347 SSRs with a relative abundance of 736.29 SSRs per Mb of the genome and with a relative density of 1249.78 bases per Mb (Table 5). Detailed distribution of the predicted SSRs is given in Table 6.

**Table 5:**
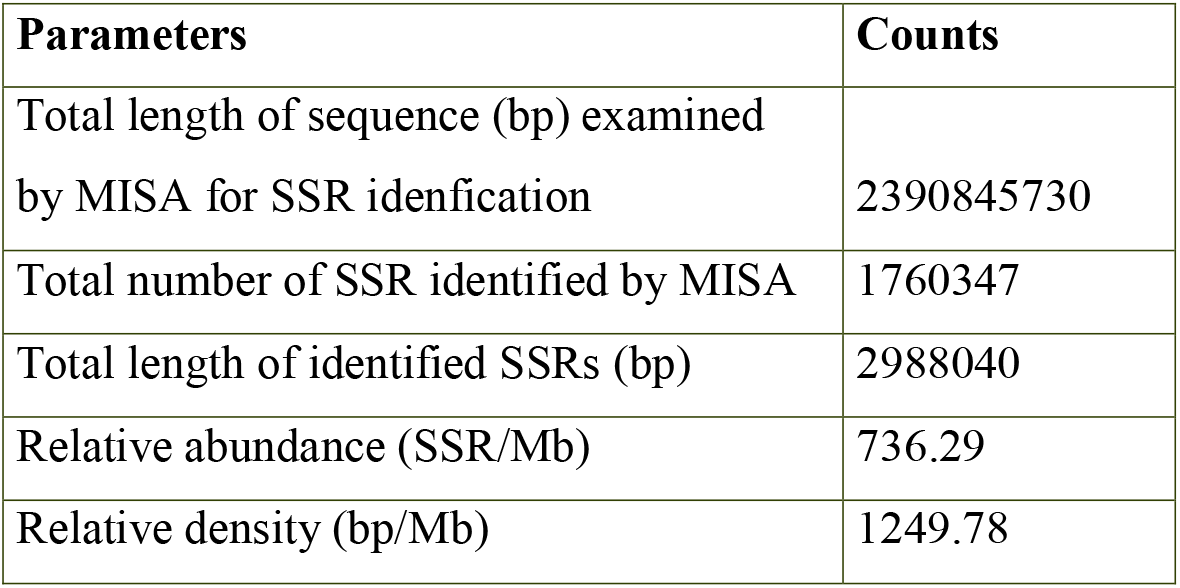
Statistics describing the SSRs identified in the Bengal tiger genome.

| Parameters | Counts |
| --- | --- |
| Total length of sequence (bp) examined by MISA for SSR identification | 2390845730 |
| Total number of SSR identified by MISA | 1760347 |
| Total length of identified SSRs (bp) | 2988040 |
| Relative abundance (SSR/Mb) | 736.29 |
| Relative density (bp/Mb) | 1249.78 |

**Table 6:**
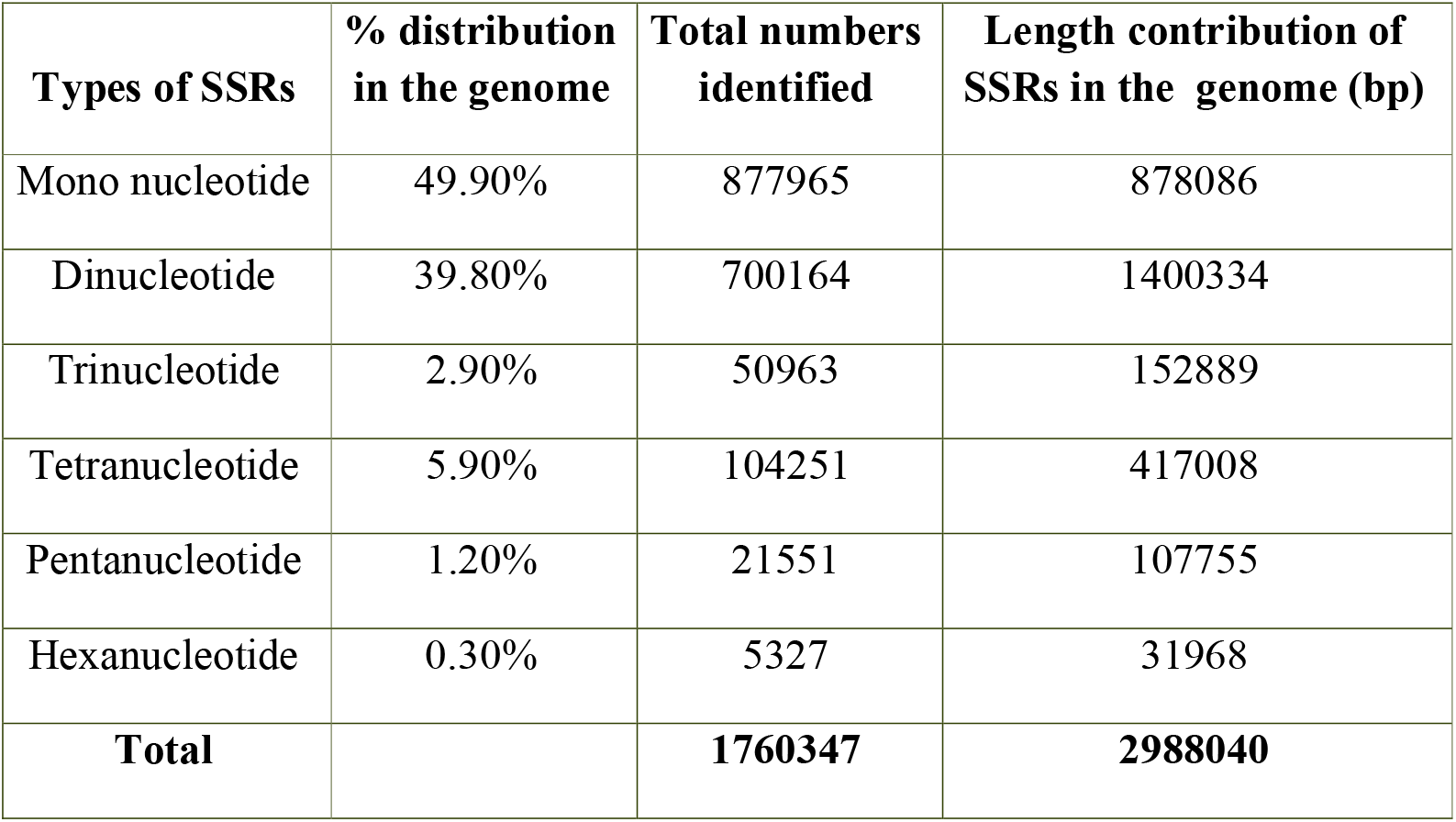
Distribution statistics of the predicted SSRs in the Bengal tiger genome.

| Types of SSRs | % distribution in the genome | Total numbers identified | Length contribution of SSRs in the genome (bp) |
| --- | --- | --- | --- |
| Mono nucleotide | 49.90% | 877965 | 878086 |
| Dinucleotide | 39.80% | 700164 | 1400334 |
| Trinucleotide | 2.90% | 50963 | 152889 |
| Tetranucleotide | 5.90% | 104251 | 417008 |
| Pentanucleotide | 1.20% | 21551 | 107755 |
| Hexanucleotide | 0.30% | 5327 | 31968 |
| **Total** |  | **1760347** | **2988040** |

## Conclusions

We report the first high coverage genome sequence of the Bengal tiger with an overview of its genomic variants when compared to the Amur tiger genome. The highlight of this work is the identification of several single nucleotide as well as large structural changes in the genome, most of which are probably individual-specific variations. However a few of these variations can potentially be associated with the animal’s ability to adapt to environmental changes, disease susceptibility and other important biological phenomena, which need to be explored further and validated with multiple individual genomes. High sequence coverage made it possible to identify copy number and other structural variants, discovery of which requires high redundancy in the genome sequence. The numerous SSRs and SNVs identified in this study can be used to strengthen forensic evidence in tiger poaching cases in India. Further as discussed by Pontius et al [31], this genome sequence and variations identified will contribute significantly to natural history studies of wild cat species which will be useful for conservation of these endangered species.

## Data availability

Mate pair library raw sequence data are deposited in NCBI with the following SRA numbers SRR5612311 and SRR5612312, while paired-end library raw sequence data are available in NCBI with the following SRA numbers SRR5591010, SRR5591011 and SRR 5591009.

## Acknowledgements

We thank Dr. Rakesh Mishra (Director, CSIR-CCMB) and Dr. K. Vasudevan for their support and encouragement. This study was funded by the Council for Industrial and Scientific Research, Govt. of India (Project code BSC0207). The authors declare no conflict of interest.

